# ZFHX3 plays a role in pain and cautious-like behaviours in mice

**DOI:** 10.1101/2024.06.18.599525

**Authors:** PM Nolan, SM Gettings, R Hillier, N Bourbia

## Abstract

A missense mutation in zinc finger homeobox-3 (ZFHX3) gene is known to alter circadian rhythms and metabolism. ZFHX3 is highly expressed in the central nucleus of amygdala and so we investigated whether female and male Zfhx3^Sci-/+^ mice have impaired pain, fear, anxiety-like, and cautious-like behaviours as these behaviours which are associated with the central nucleus of the amygdala. Using mechanical and thermal sensitivity test, the fear conditioning test and the light-dark box test, we found that female Zfhx3^Sci-/+^ mice have hypoalgesia to mechanical stimulus while hyperalgesia to heat stimulus. Additionally, both female and male Zfhx3^Sci-/+^ display reduction of cautious-like behaviour. While neither female and male Zfhx3^Sci-/+^ mice showed a difference in fear conditioning or anxiety-like behaviour, we have demonstrated for the first time that ZFHX3 is involved in pain and cautious behaviours.

**Highlights:**

- Female and male Zfhx3^Sci/+^ mice displayed less cautious-like behaviour
- Zfhx3^Sci/+^ female mice had hypoalgesia to noxious mechanical stimulus
- Zfhx3^Sci/+^ female mice had hyperalgesia to noxious heat stimulus

## Introduction

ZFHX3, also known as Zinc Finger Homeobox 3 or ATBF1, is a transcription factor binding an AT (adenine and thymine)-rich motif (Morinaga et al., 1991; Yasuda et al., 1994). The role of ZFHX3 has been well characterised in circadian rhythms (Balzani et al., 2016; Hughes et al., 2021; Parsons et al., 2015; Wilcox et al., 2017, 2021), as well as metabolism (Nolan et al., 2023; Park, 2022) as ZFHX3 is present in the suprachiasmatic nucleus and the hypothalamus respectively. ZFHX3 also play a role in neurodevelopment (Pérez Baca et al., 2024), and causes spinocerebellar ataxia (Figueroa et al., 2024; Wallenius et al., 2024) and epilepsy (He et al., 2024). This shows that ZFHX3 is involved in various neurophysiology and neuropathology aspects.

However, the role of ZFHX3 in pain and nociception, anxiety, and fear are either poorly understood or has not been investigated, despite ZHFX3 being highly expressed in the central nucleus of amygdala (Hochgerner et al., 2023; Lein et al., 2007; Yeh et al., 2024).

The central nucleus of amygdala is well known to play a key role in sensory-aspect and emotional pain (Ansah et al., 2010; Bourbia et al., 2010), fear (LeDoux, 2003), and anxiety (Fox & Shackman, 2019; McNaughton & Corr, 2004). Therefore, male and female Zfhx3^Sci/+^ mice (developed by Parsons et al., 2015, and hosted in the Mary Lyon Center) where selected as a model to investigate whether a point mutation in ZFHX3 induces changes in anxiety-like and cautious-like behaviours (using the light-dark box test), pain and nociception (through assessing reflex withdrawal to a mechanical and thermal stimulus), and fear behaviours (using fear conditioning) associated to the central nucleus of amygdala.

## Materials and methods

### Mice

All animal work was performed under the guidance issued by the Medical Research Council and Home Office Project License 30/3206, with local ethical approval.

Zfhx3^Sci/+^ mutant mice were bred in Mary Lyon Centre at the MRC Harwell Institute. The original mutation, found and described previously (Parsons et al., 2015), was initially bred in both C57BL/6J and C3H/HeH backgrounds before being maintained in a mixed C3H/HeH × C57BL6/J F1 background. A homozygote is fatal in this mouse line, as such this study used heterozygote mice. Mice will be referred in this publication including in the graphs as female or male SCI wild type (WT) for Zfhx3^+/+^ or SCI heterozygote (HET) for Zfhx3^Sci-/+^.

Male and female SCI were used and kept by sex in mixed-genotype groups in a 12-hour light-dark cycle (lights on at 07:00; lights off at 19:00) with *ad libitum* food (RM3 food) and water.

The mice were housed by group of 4-5 in Blue Line IVC cages (Techniplast) (base size: 35 × 14 × 19cm LxHxW), containing Eco-Pure Lab Animal Bedding (Datesand) and FDA Paper Shavings (Datesand).

Tunnel or cupping methods of handling were used for routine animal checks and cage changes, while the tail handling method was used to transfer animals to the experiment tasks.

### Behavioural testing

All behavioural tests were randomized and performed blind to the experimenter, and they were also described previously (Bourbia et al., 2019). Behavioural testing was performed between 8 to 12 weeks of age. Mechanical and thermal sensitivity tests, and light-dark box were performed between 13:00 and 17:00, and fear-conditioning test was performed between 10:00 and 12:00 during the first day of the protocol and between 10:00 and 18:00 during the second day of the protocol.

### Mechanical (von Frey filament test) and thermal Sensitivity (hot plate test)

Mechanical sensitivity was assessed by measuring the minimum force needed to induce a hind limb withdrawal response to a mechanical stimulus induced by calibrated monofilament from a force of 0.8 to 10 g (Touch Test™ Sensory Evaluator, Stoelting Europe, Ireland) applied under the foot pad. The mice were habituated to the experimental condition for 1 hour on 2 days followed by testing on day 3. The lowest force to induce 100 % withdrawal response based on five stimuli per force was considered as the withdrawal threshold force expressed in grams (g).

Thermal sensitivity was assessed using the hot plate test set (Bioseb, Chaville, France), at 52 °C, where mice were placed on top until the first paw licking is observed. Mice were then immediately removed from the equipment and returned to their home cage. The latency to the first paw licking, expressed in seconds was considered as the withdrawal latency.

### Fear conditioning

Pavlovian threat conditioning was assessed using the Fear Conditioning System (Ugo Basile, Italy). The Fear conditioning consists of 3 phases, where mice are returned to their home-cage after each phase:

- Conditioning phase: mice were placed in a square chamber with an electric grid on the floor for 10 min, where mice are exposed to a series of three conditioned stimuli (5 s tone) each coupled with a 0.5 s footshock stimulus (0.5 mA) at 150 s, 305.5 s and 461 s.
- Context phase: 24 h after conditioning, mice were placed in the same chamber and freezing behaviour recorded for 5 min in the absence of any stimuli.
- Cue phase: 4 h after the context phase, mice were placed in a round Plexiglas cylinder with vanilla fragrance spread at the top of the cylinder for 6 min during which the mouse is exposed to the same sound as in the conditioning phase.

Freezing time expressed in seconds was measured using the ANYMaze video tracking system (Stoelting Europe).

### Light dark box

A 40 cm × 40 cm × 40 cm plastic box, equally divided into dark (0 lux) and light (100 lux) areas with free access between areas through a 3 cm x 3 cm open door, was used to assess anxiety-like behaviour. The mice were placed inside the dark or light area and were able to explore both over a 5 min period. The time spent in each area, the number of entries, and the latency to the first entry in the light area (only when mice were initial placed in the dark area) were measured via video tracking using EthoVision XT software (Noldus, Wageningen, The Netherlands) and number of cautious-like and non-cautious-like events were measured manually by the experimenter. Two types of cautious behaviour and one type of non-cautious behaviour were recorded:

- Cautious behaviour: Informed exploration when the mice in the dark area pass the head through the open door and pause 2 seconds before continuing entering the light area (Supplemental Video 1).
- Cautious behaviour: Informed avoidance when the mice in the dark area pass the head through the open door followed by retracting their head and choosing to remain in the dark area (Supplemental Video 2).
- Non-cautious behaviour was observed when the mice in the dark area walk straight through the door into the light area without pausing (Supplemental Video 3).

### Statistical analysis

Statistical analysis was performed using GraphPad Prism 9.4.0. 2-way ANOVA followed by a Tukey’s multiple comparisons test were used for multiple comparison. Unpaired t-test, Mann-Whitney or Welch’s t test, based on Shapiro-Wilk test for normality, were used to compare genotypes per sex group. All figures were formatted in Inkscape V.1.1.2.

## Results

### Mechanical sensitivity

Mechanical stimuli to the foot pad inducing hind limb withdrawal reflexes were used to assess whether a point mutation in ZHFX3 would induce mechanical allodynia, hyper- or hypoalgesia.

Female SCI HET has a significant increase of the mechanical threshold compared to female SCI WT in both left and right paw (2-way ANOVA, *N* = 7 to 9; Genotype: *P* < 0.0001, *F*_1,28_ = 21.07, Paw: *P* = 0.3178, *F*_1,28_ = 1.034, Interaction: *P* = 0.6707, *F*_1,28_ = 0.6707, Figure 2A) suggesting mechanical hypoalgesia, while there is no difference between genotypes or paw sensitivity in male SCI (2-way ANOVA, *N* = 6 to 8; Genotype: *P* = 0.5682, *F*_1,24_ = 0.3348, Paw: *P* = 0.7331, *F*_1,24_ = 0.1190, Interaction: *P* = 0.7004, *F*_1,24_ = 0.1517, Figure 2B).

**Figure 1:**
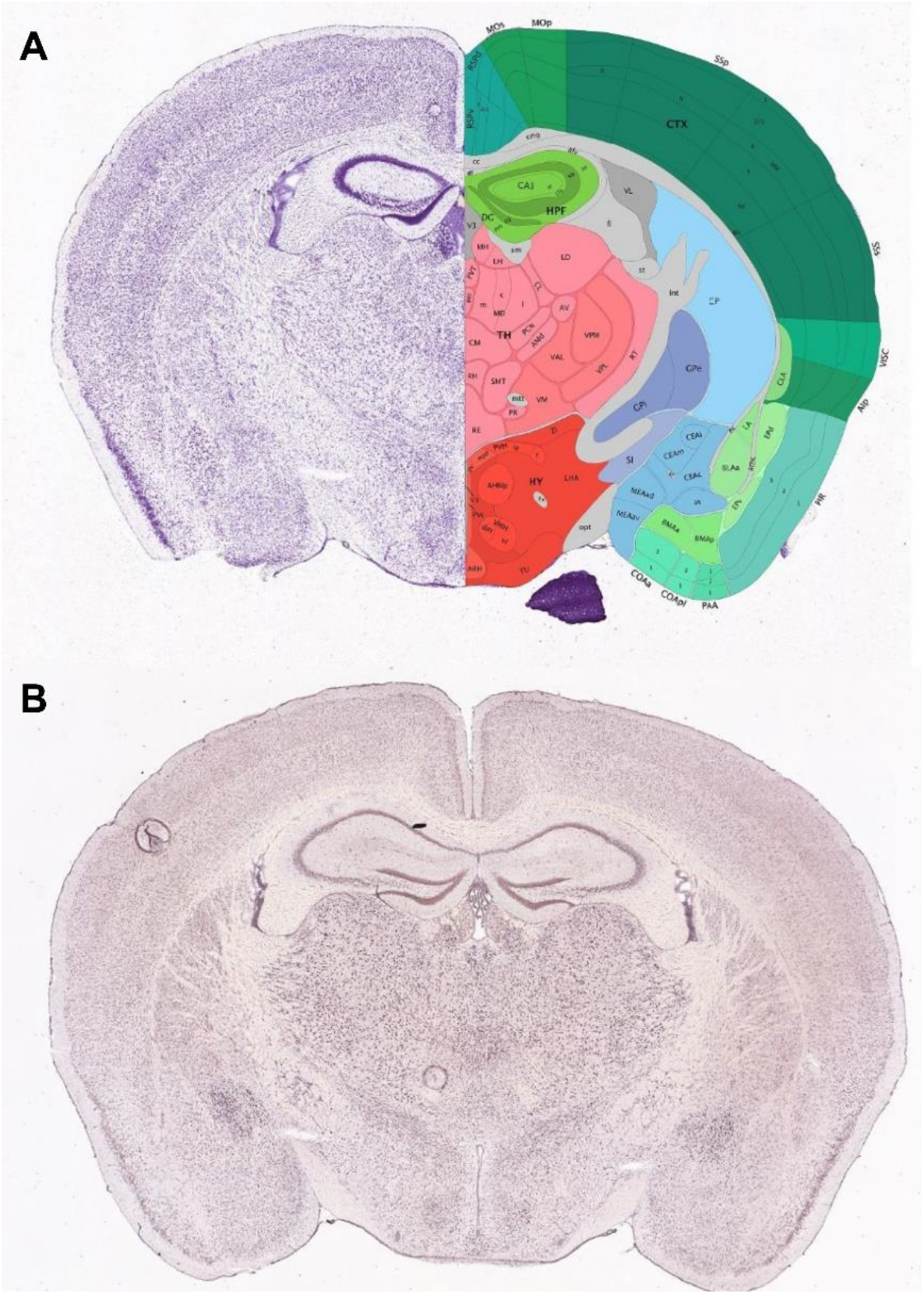
ZFHX3 expression in adult mouse brain. A: Nissl (left) and anatomical annotations (right) from the Allen Mouse Brain Atlas and Allen Reference Atlas – Mouse Brain, at the same slice position as B. Allen Mouse Brain Atlas, mouse.brain-map.org and atlas.brain-map.org. B: Expression of ZFHX3 in adult mouse brain. Allen Mouse Brain Atlas, https://mouse.brain-map.org/experiment/show/74641308.

**Figure 2.**
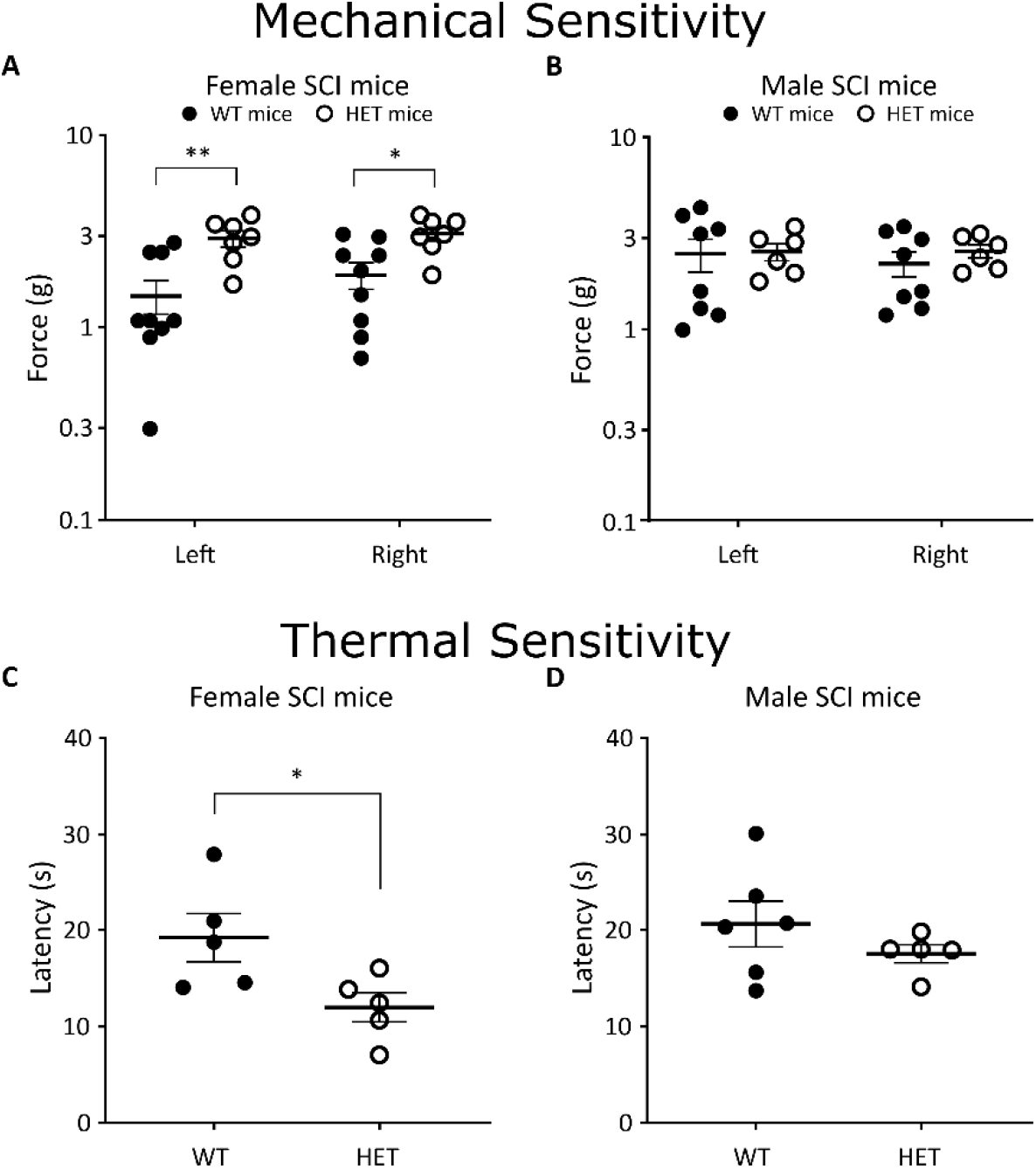
Mechanical and thermal sensitivity in SCI WT (black round) and SCI HET (white round) mice. The mechanical sensitivity of female (A) and male (B) mice was measured as withdrawal reflexed of the hind paw to mechanical stimulus force expressed in g. Thermal sensitivity in female (C) and male (D) measured as the latency to observe a reflex response (lick of the paw) induced by hot plate temperature. * P ≤ 0.05, ** P ≤ 0.01. WT = wild type, HET = heterozygote.

### Thermal sensitivity

Thermal thresholds inducing hind limb withdrawal licking reflexes were used to assess whether a point mutation in ZHFX3 would induce thermal allodynia, hyper- or hypoalgesia. Female SCI HET has a significant increase to thermal sensitivity compared to female SCI WT (*P* = 0.0317, *U* = 2, *N* = 5, Figure 2C) suggesting thermal allodynia, while there is no difference between genotypes of male SCI (*P* = 0.3052, *U* = 9, *N* = 6 and 5, Figure 2D)

### Fear Conditioning

During the 3 phases of the fear conditioning, male and female mice had an increased freezing time associated with conditioning (Figure 3), context and cue (Figure 4). However, there was no difference between SCI WT and SCI HET in both male and female mice.

**Figure 3.**
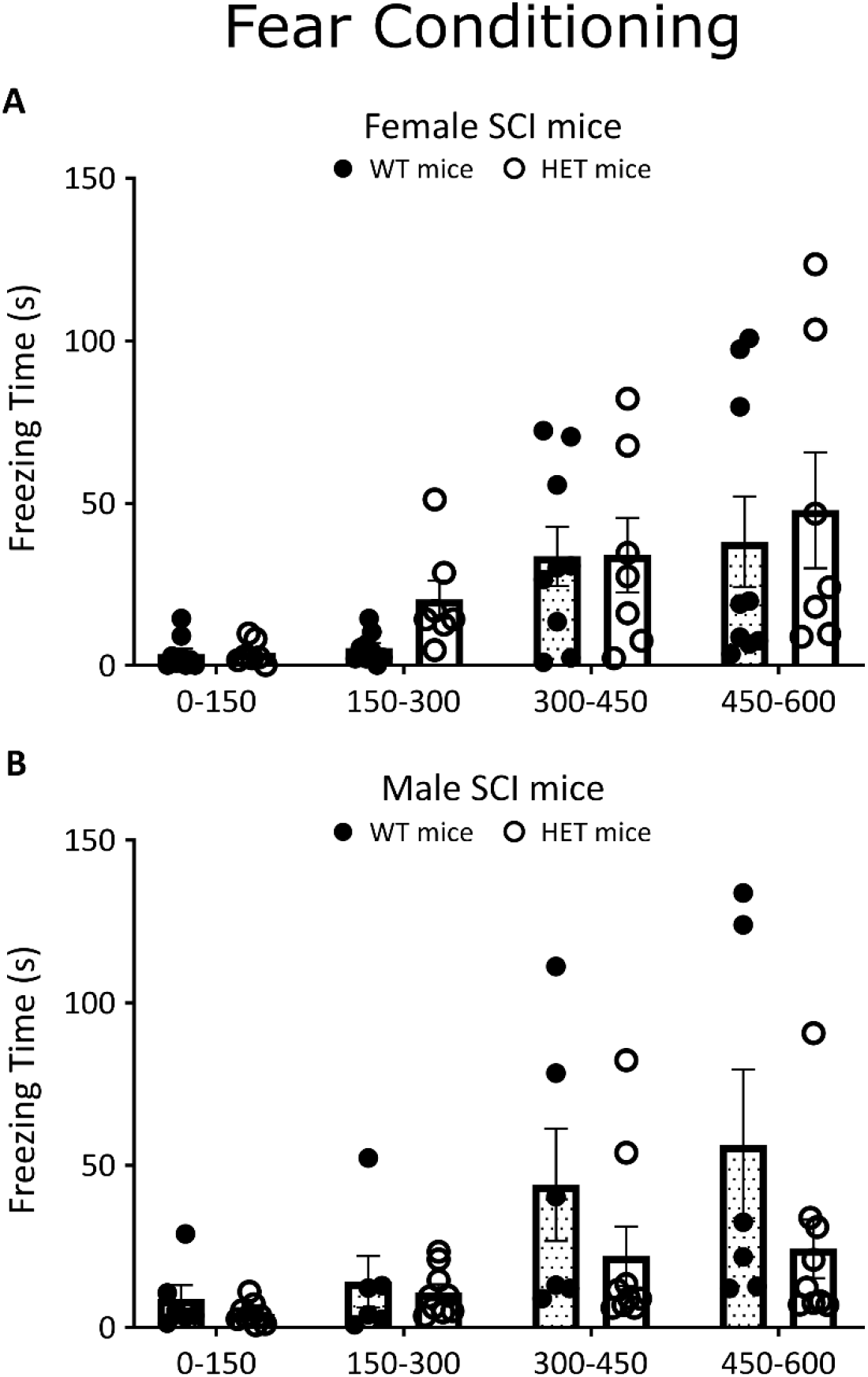
Fear Conditioning of SCI WT (black round) and SCI HET (white round) mice. Freezing time per 150 sec intervals expressed in seconds during the conditioning phase of the Fear Conditioning test in female (A) and male (B) mice. WT = wild type, HET = heterozygote.

**Figure 4.**
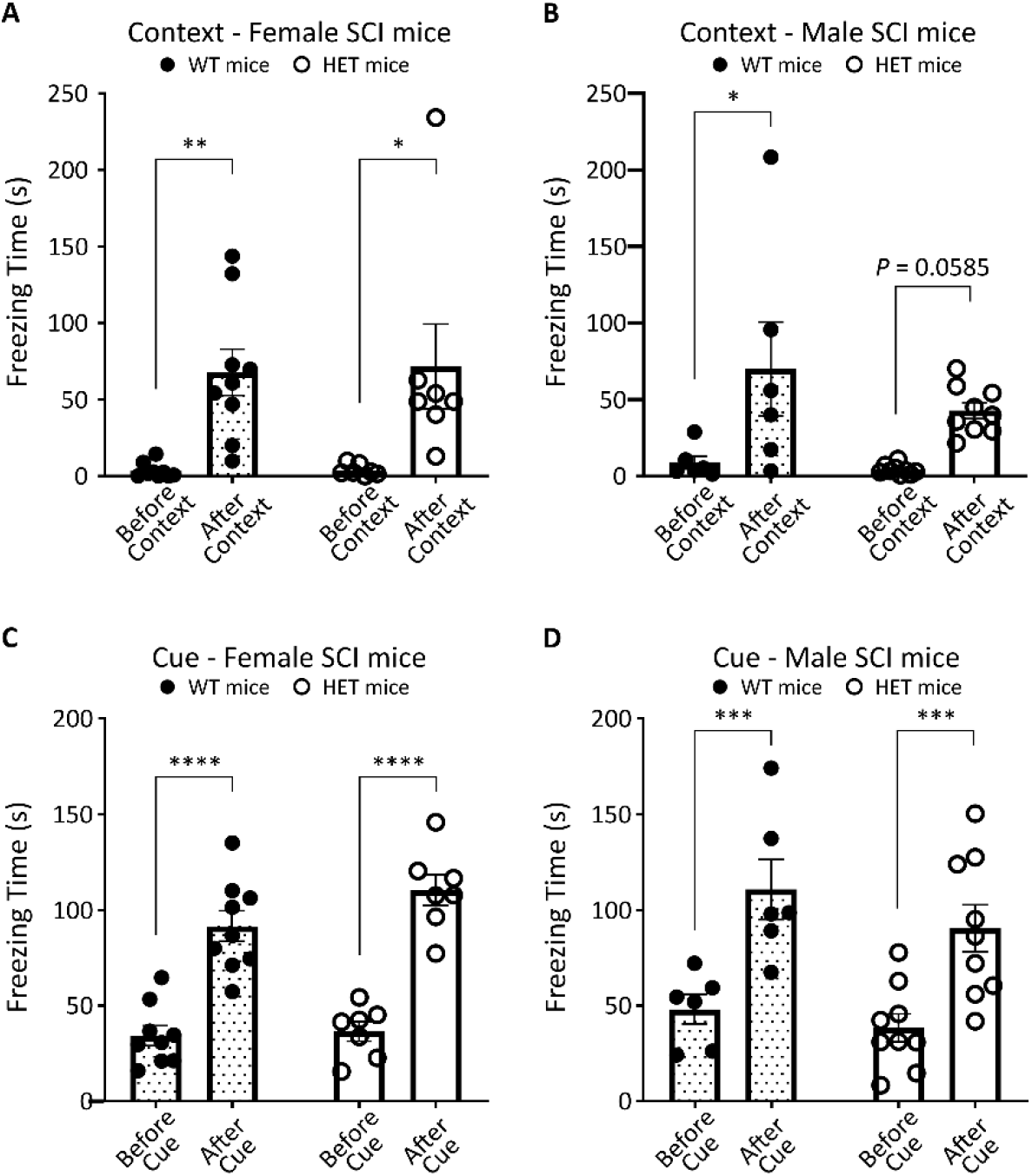
Cue and context fear conditioning of SCI WT (black round and dotted bar) and SCI HET (white round and white bar) mice. Freezing time expressed in second during the context (A,B) and Cue (C,D) phase of the Fear Conditioning test in female (A, C) and male (B, D) mice. * P ≤ 0.01, ** P ≤ 0.01, *** P ≤ 0.001, **** P ≤ 0.0001. WT = wild type, HET = heterozygote.

During the conditioning phase, both female (2-way ANOVA, *N* = 9 and 7; Genotype: *P* = 0.5538, *F*_1,14_ = 0.3680; Freezing time every 150s: *P* = 0.0014, *F*_1.161,16.26_ = 13.56; mouse variation: *P* < 0.001, *F*_14,42_ = 4.526; Interaction: *P* = 0.6586, *F*_3,42_ = 0.5384, Figure 3A) and male (2-way ANOVA, *N* = 6 and 9; Genotype: *P* = 0.1877, *F*_13, 39_ = 1.934; Freezing time every 150s: *P* = 0.0057, *F*_1.321,17.18_ = 8.660; mouse variation: *P* = 0.003, *F*_13, 39_ = 4.114; Interaction: *P* = 0.2011, *F*_3,39_ = 1.617, Figure 3B) mice had an increase of freezing time induced by footshock.

During the context phase, female (2-way ANOVA, *N* = 9 and 7; Genotype: *P* = 0.8906, *F*_1,14_ = 0.8906; Freezing time every 150s: *P* = 0.0005, *F*_1,14_ = 20.58; mouse variation: *P* = 0.4431, *F*_14,14_ = 1.081; Interaction: *P* = 0.9063, *F*_1,14_ = 0.01437, Figure 4A) and male (2-way ANOVA, *N* = 6 and 9; Genotype: *P* = 0.2449, *F*_1,13_ = 1.484; Freezing time every 150s: *P* = 0.0015, *F*_1,13_ = 15.98; mouse variation: *P* = 0.4322, *F*_13,13_ = 1.101; Interaction: *P* = 0.3794, *F*_1,13_ = 0.8279, Figure 4B) mice had an increased freezing time during the context time compared to the 150 s pre condition period measured during the conditioning phase.

During the cue phase, female (2-way ANOVA, *N* = 9 and 7; Genotype: *P* = 0.2155, *F*_1,14_ = 1.683; Freezing time every 150s: *P* < 0.0001, *F*_1,14_ = 154.9; mouse variation: *P* = 0.0561, *F*_14,14_ = 2.405; Interaction: *P* = 0.1362, *F*_1,14_ = 2.499, Figure 4C) and male (2-way ANOVA, *N* = 6 and 9; Genotype: *P* = 0.3024, *F*_1,13_ = 1.153; Freezing time every 150s: *P* < 0.0001, *F*_1,13_ = 55.73; mouse variation: *P* = 0.0193, *F*_13,13_ = 3.328; Interaction: *P* = 0.5010, *F*_1,13_ = 0.5010, Figure 4D) mice had an increased freezing time during the cue time compared to the pre-cue time.

### Anxiety-like behaviour

The light-dark box was used to assess anxiety-like and cautious-like behaviour. Two protocols were used where mice were first placed in the dark or the light areas.

#### Mouse starting in the dark area

We confirmed that female (2-way ANOVA, *N* = 5 and 4; Light/Dark area effect: *P* < 0.0001, *F*_1,14_ = 43.66; Genotype: *P* = 0.9653, *F*_1,14_ = 0.001962; Interaction: *P* = 0.3109, *F*_1,14_ = 1.105, Figure 5A) and male (2-way ANOVA, *N* = 5 and 5; Light/Dark area effect: *P* < 0.0001, *F*_1,16_ = 29.04; Genotype: *P* = 0.9811, *F*_1,16_ = 0.0005764; Interaction: *P* = 0.4638, *F*_1,16_ = 0.5634, Figure 5B) mice spent more time in the dark area than the light area, independently of their genotype.

**Figure 5.**
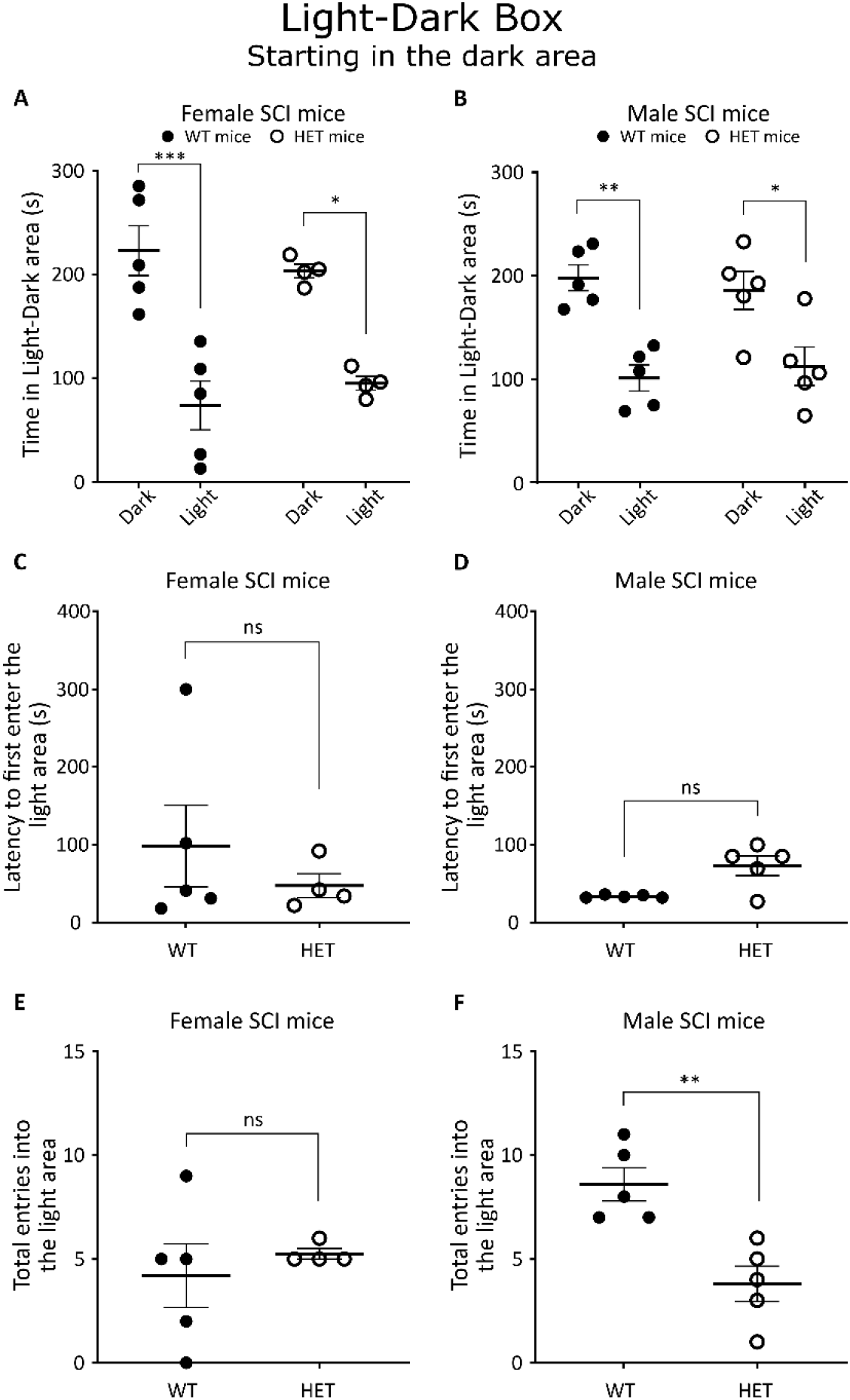
The behaviour of female (A, C, E) and male (B, D, F) SCI WT (black round) and HET (white round) mice starting in the dark area of the Light-Dark Box test. A Time spend in each area (A and B) and latency to the first entry in the light area (C and D) are expressed in second. E and F represent the number of entries in the light area. * P ≤ 0.01, ** P ≤ 0.01, ns = P > 0.05. WT = wild type, HET = heterozygote.

Male SCI HET mice had an increased latency to go into the light area compared to Male SCI WT (*P* = 0.0341, *t* = 3.149, *df* = 4.03, Figure 5D), while female SCI do not show difference in latency to enter the light area between both genotypes (*P* = 0.9048, *U* = 9, Figure 5C). One female SCI WT never entered in the light area.

Additionally male SCI HET mice had a reduction of number of entries in the light area compared to the SCI WT mice (*P* = 0.0036, *t* = 4.057, *df* = 8, Figure 5F), whereas there was no difference in female SCI genotypes (*P* = 0.6349, *U* = 7, Figure 5E).

#### Mouse starting in the light area

We confirmed that female mice independently of their genotype spent more time in the dark area (2-way ANOVA, *N* = 9 and 7; Light/Dark area effect: *P* < 0.0001, *F_1,28_* = 31.87; Genotype: *P* = 0.9974, *F*_1,28_ = 1.123e^-005^; Interaction: *P* = 0.5690, *F*_1,28_ = 0.3321, Figure 6A). The male mice spent more time in the dark area, however there was an interaction difference despite no genotype difference (2-way ANOVA, *N* = 7 and 7; Light/Dark area effect: *P* = 0.0002, *F_1,24_* = 19.11; Genotype: *P* = 0.9952, *F*_1,24_ = 3.675e^-005^; Interaction: *P* < 0.0001, *F*_1,24_ = 23.84, Figure 6B). Additionally, Tukey’s multiple comparison showed no difference between time spent in the light and dark area in male SCI HET (*P* = 9834, Dark area: mean time = 145.6s, *SEM* = 20.2; Light area: mean time = 153.4s, *SEM* = 20.2).

**Figure 6.**
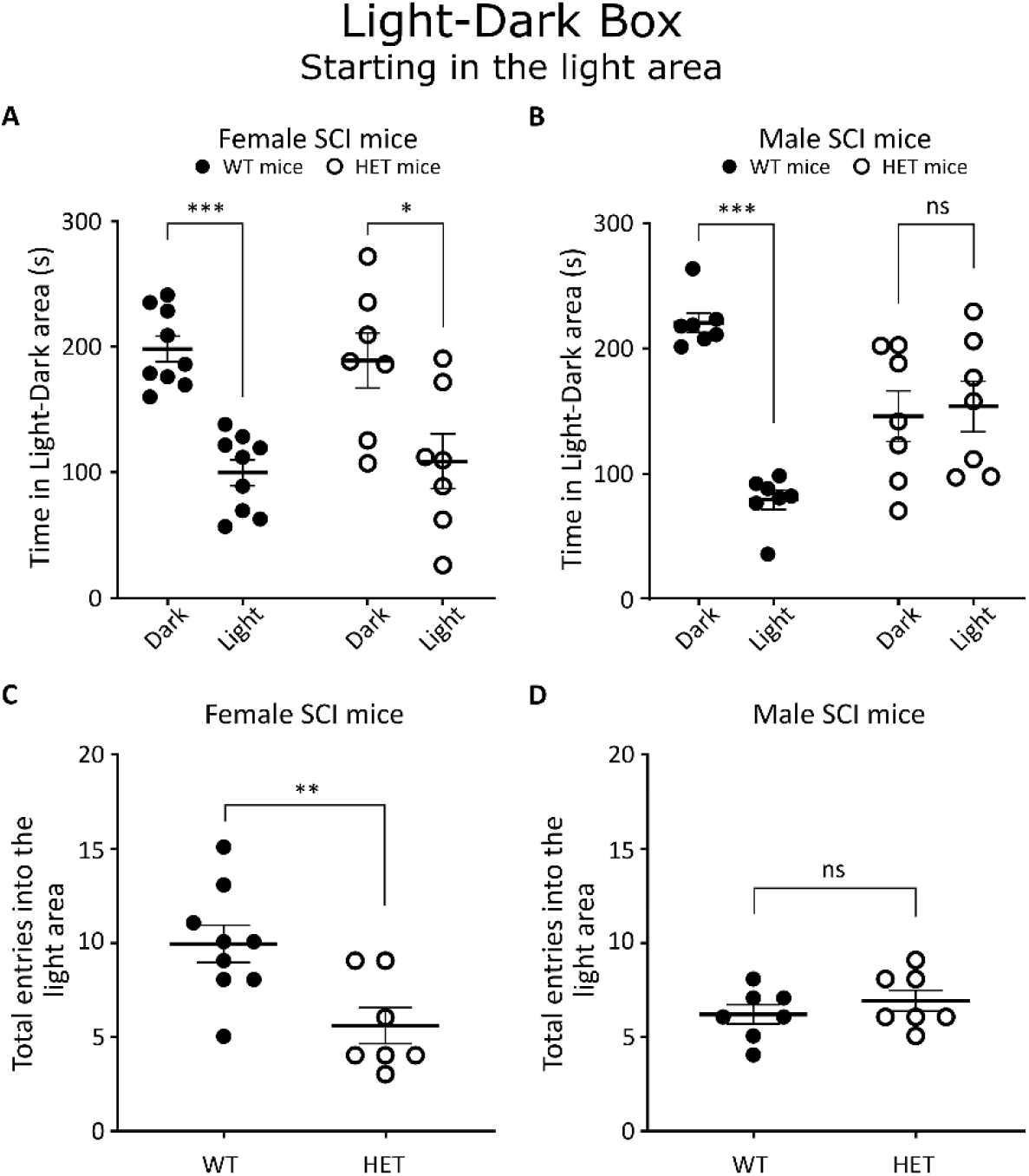
The behaviour of female (A, and C) and male (B and D) SCI WT (black round) and HET (white round) mice starting in the light area of the Light-Dark Box test. Time spent in each area (A and B) is expressed in second. C and D represent the total of entries in the light area. * P ≤ 0.01, ** P ≤ 0.01, *** P ≤ 0.001, ns = P > 0.05. WT = wild type, HET = heterozygote.

Female SCI HET mice had a reduction of number of entries in the light area compared to female SCI WT (*P* = 0.0078, *t* = 3.105, *df* = 14, Figure 6C), whereas there was no difference between male SCI genotypes (*P* = 0.3606, *t* = 0.9506, *df* = 12, Figure 6D).

### Cautious-like behaviour

The light-dark box was used to assess non-cautious-like and both cautious-like (informed exploration and informed avoidance) behaviour in SCI mice.

#### Mouse starting in the dark area

Female SCI HET only demonstrated a reduction in one aspect of cautious-like behaviour, the informed avoidance (*P* = 0.0286, *U* = 0, Figure 7C), but do not show a significant difference in the informed exploration (*P* = 0.1143, *U* = 2.5, Figure 7A) nor non-cautious-like behaviour (*P* = 0.1405, *t* = 1.698, *df* = 6, Figure 7E) when compared to female SCI WT.

**Figure 7.**
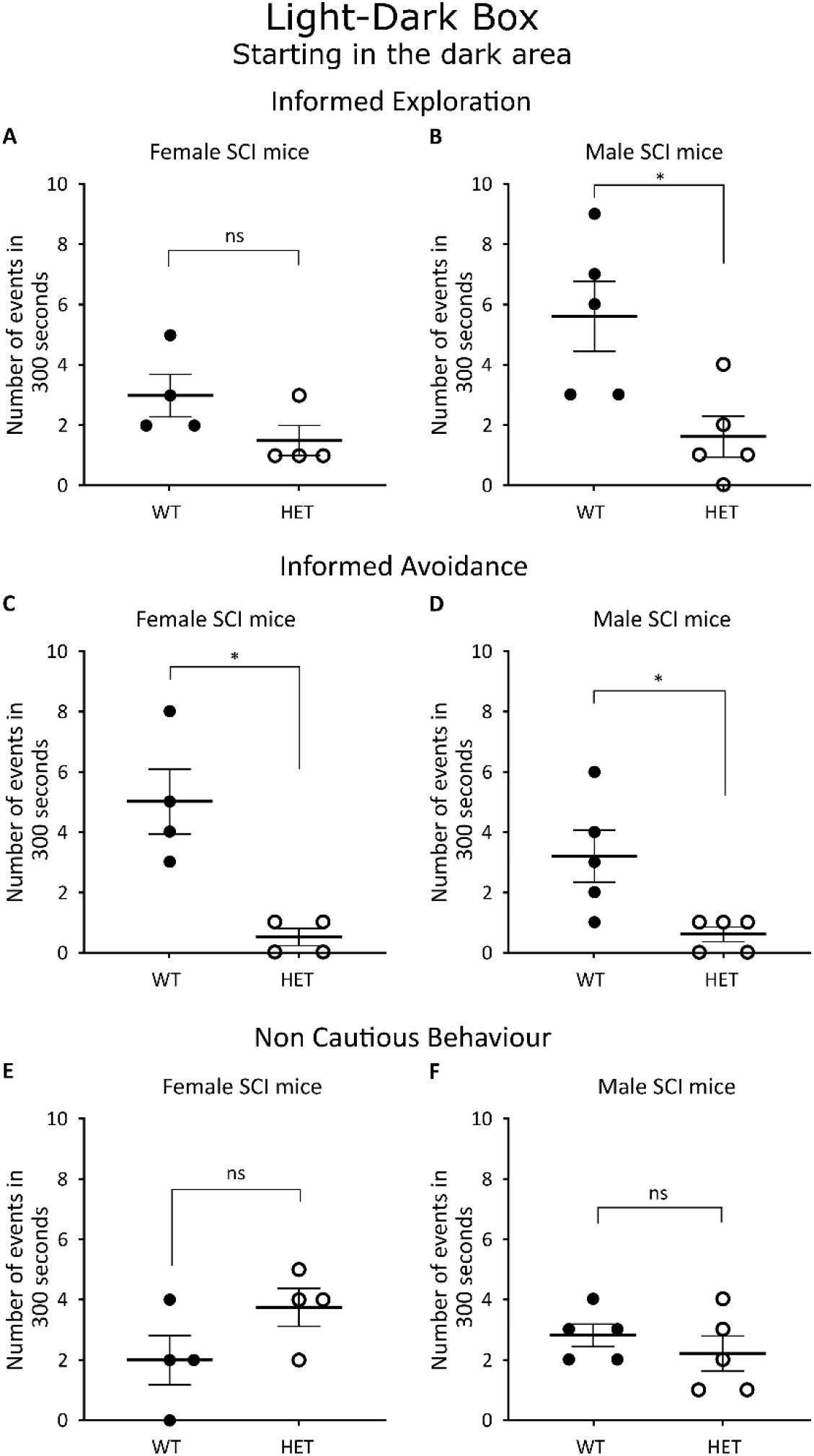
Cautious and non-cautious behaviour when female (A, C and E) and male (C, F and F) SCI WT (black round) and HET (white round) mice start in the dark area of the Light-Dark Box. Number of informed exploration (A and B), informed avoidance (C and D) and non-cautious behaviour (E and F) is recorded within the 300s of the light-dark box test. * P ≤ 0.01, ns = P > 0.05. WT = wild type, HET = heterozygote. WT = wild type, HET = heterozygote.

Male SCI HET showed a reduction of both cautious-like behaviour, informed exploration (*P* = 0.0180, *t* = 2.965, *df* = 8, Figure 7B) and informed avoidance (*P* = 0.0317, *U* = 1.5, Figure 7D), but showed no difference in non-cautious-like behaviour compared to male SCI WT (*P* = 0.4117, *t* = 0866, *df* = 8, Figure 7F).

#### Mouse starting in the light area

Female SCI HET exhibited a reduction of both cautious-like behaviour, informed exploration (*P* < 0.0001, *t* = 6.212, *df* = 14, Figure 8A) and informed avoidance (*P* = 0.0003, *U* = 1, Figure 8C) compared to female SCI WT, but do not show changed in non-cautious-like behaviour (*P* = 0.0663, *U* = 14.5, Figure 8E).

**Figure 8.**
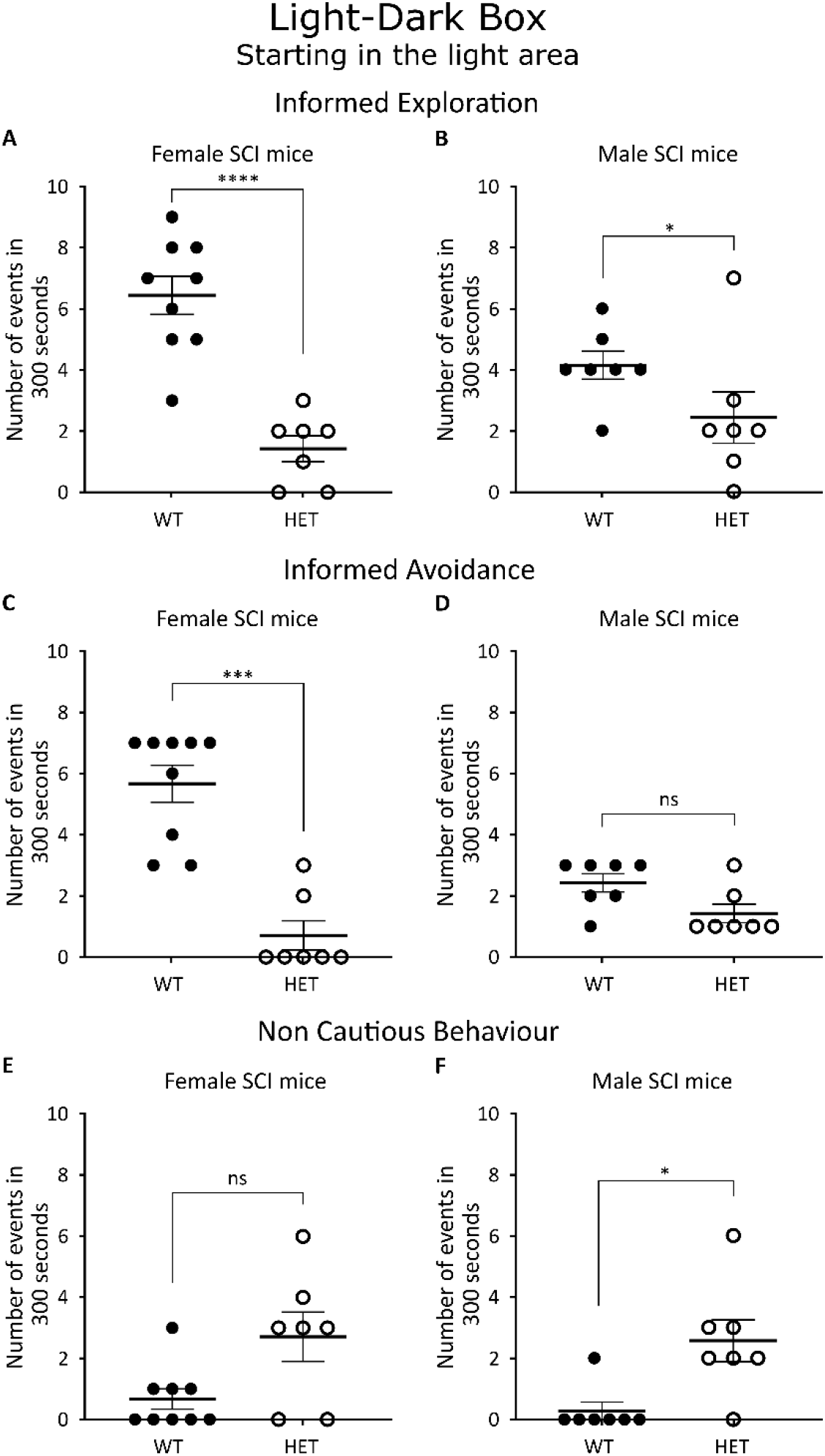
Cautious and non-cautious behaviour when female (A, C and E) and male (C, F and F) SCI WT (black round) and HET (white round) mice start in the light area of the Light-Dark Box. Number of informed exploration (A and B), informed avoidance (C and D) and non-cautious behaviour (E and F) is recorded within the 300s of the light-dark box test. * P ≤ 0.01, *** P ≤ 0.001, **** P ≤ 0.0001 ns = P > 0.05. WT = wild type, HET = heterozygote.

Male SCI HET only displayed an increase of non-cautious-like behaviour compared to male SCI WT (*P* = 0.0169, *U* = 5.5, Figure 8 B), but demonstrated no difference in both informed exploration (*P* = 0.0989, *t* = 1.789, *df* = 12, Figure 8D) and informed avoidance (*P* = 0.0676, *U* = 9.5, Figure 8F).

## Discussion

The same point of mutation in ZFHX3 gene that induces circadian rhythms and metabolism dysfunctions also affects pain and cautious-like behaviours, functionally link to the central nucleus amygdala where ZFHX3 is highly expressed.

ZFHX3 does not seem to be involved in fear conditioning nor anxiety-like behaviour in male and female mice. However, ZFHX3 is involved in cautious behaviour and both female and male Zfhx3^Sci/+^ showed a reduction in cautious behaviour, despite some difference associated to the light-dark box experimental design. The initial placement of mouse in the light dark box (starting from the dark or lite area) is known to affect some outcomes of the light-dark box test (Campos-Cardoso et al., 2023; Kulesskaya & Voikar, 2014).

As defined in the study by Zhou et al. 2022, caution is taking care to avoid errors or danger (Gold & Shadlen, 2007; Zhou et al., 2022). The study conducted by Zhou et al., (2022) has also shown that cautious behaviour is not caused by anxiety, and doesn’t require the frontal cortex as proposed by other studies (Maanen et al., 2011; Meyer & Bucci, 2016). We know that central amygdala is involved in avoidance behaviour (Grossman et al., 1975), defensive avoidance and defensive approach (McNaughton & Corr, 2004), and in sensing environmental setting for decision of action making (Isa et al., 2022).

Therefore, it is possible to hypothesize that a ZFHX3-positive neuron population inside the central nucleus amygdala could be linked to cautious behaviour. Using conditional ZFHX3 (in central amygdala) KO mice, in addition to further studies, could help to answer this hypothesis. ZFHX3 is a transcription factor, and it would be interesting to understand the molecular mechanism underlying cautious behaviour. For instance, this same Zfhx3^Sci/+^ mouse line has a decrease of somatostatin and tachykinin 2 in the suprachiasmatic nucleus (Parsons et al., 2015), and somatostatin and tachykinin 2 are associated to fear conditioning (McCullough et al., 2018) yet we did not observed difference in fear conditioning between the SCI WT and HET mice. Therefore, while there is a knowledge gap on the circuitry of cautious behaviour, it likely involves ZFHX3.

Mice with a knockout of ZFHX2 have hypoalgesia to noxious mechanical stimulus but have hyperalgesia to noxious heat (Habib et al., 2018). This pain and nociception pattern is similar to the one observed in our study, however we only observed it in female mice SCI HET not in male mice. ZFHX2 is an homologous of ZFHX3 and also expressed in the DRG neuron (Megat et al., 2019). Therefore, while we chose to study pain-related behaviour because ZFHX3 is highly express in CeA, it is possible that the pain difference in female SCI HET is caused by changes in the dorsal root ganglia and nociceptors in addition or instead of amygdaloid ZFHX3. Especially that somatostatin is expressed in DRG neuron and lack of DRG somatostatin induces an increase sensitivity to noxious heat and to mechanical stimuli (Huang et al., 2018).

Sex difference in pain and nociception is known and involved cellular and molecular differences in pain processing that are sex specific (Presto et al., 2022; Sorge & Totsch, 2017). As ZFHX3 is a transcription factor, its mutation may affect the expression of a key protein predominantly involved in female pain processing that needs to be determined with further studies, as well as to understand whether ZFHX3 modulates the central or peripheral pain and nociception system in female mice.

In conclusion, this behavioural study has shown that ZFHX3 is involved in cautious behaviour in male and female mice, and in pain and nociception in female but not male mice. Further studies are necessary to identify the neuroanatomic functions associated to ZFHX3 but also the neurogenomic aspect of ZFHX3 as it is a transcription factor in the context of pain and nociception.

## CRediT authorship contribution statement

**Patrick M Nolan**: Resources, Funding acquisition, Supervision, Writing – review & editing. **Sean M Gettings**: Visualization, Writing – review & editing. **Rosie Hiller**: Writing – review & editing. **Nora Bourbia**: Conceptualization, Data curation, Formal Analysis, Investigation, Methodology, Project administration, Validation, Writing – original draft.

## Acknowledgments

This research project was supported by the Medical Research Council (MC_U142684173). We thank the Mary Lyon Center for their support in hosting and caring for our mice.

